# Fc-engineered antibodies leverage neutrophils to drive control of *Mycobacterium tuberculosis*

**DOI:** 10.1101/2022.05.01.490220

**Authors:** Edward B. Irvine, Joshua M. Peters, Richard Lu, Patricia S. Grace, Jaimie Sixsmith, Aaron Wallace, Matthew Schneider, Sally Shin, Wiktor Karpinski, Jeff C. Hsiao, Esther van Woudenbergh, Arturo Casadevall, Bryan D. Bryson, Lisa Cavacini, Galit Alter, Sarah M. Fortune

**Affiliations:** Ragon Institute of MGH, MIT and Harvard, Cambridge, MA, USA; Department of Immunology and Infectious Diseases, Harvard T.H. Chan School of Public Health, Boston, MA, USA; Department of Biological Engineering, Massachusetts Institute of Technology, Cambridge, MA, USA; MassBiologics of the University of Massachusetts Chan Medical School, Boston, MA, USA; Department of Molecular Microbiology and Immunology, Johns Hopkins Bloomberg School of Public Health, Baltimore, MD, USA; Division of Infectious Disease, Massachusetts General Hospital, Boston, MA, USA

## Abstract

Novel vaccination and therapeutic strategies are urgently needed to mitigate the tuberculosis (TB) epidemic. While extensive efforts have focused on potentiating cell-mediated immunity to control *Mycobacterium tuberculosis* (*Mtb*) infection, less effort has been invested in exploiting the humoral immune system to combat *Mtb*. Emerging data point to a role for antibodies in microbial control of *Mtb*, however the precise mechanism(s) of this control remain incompletely understood. Here we took an antibody Fc-engineering approach to determine whether Fc-modifications could improve the ability of antibodies to restrict *Mtb*, and to define Fc-mediated mechanism(s) antibodies leverage for this restriction. Using an antibody specific to the capsular polysaccharide α-glucan, we engineer a panel of Fc variants to augment or dampen select antibody effector functions, rationally building antibodies with enhanced capacity to promote *Mtb* restriction in a human whole blood model of infection. Surprisingly, restrictive Fc-engineered antibodies drive *Mtb* control in a neutrophil, not monocyte, dependent manner. Using single cell RNA sequencing, we show that restrictive antibodies promote neutrophil survival and expression of cell intrinsic antimicrobial programs. These data provide a roadmap for exploiting Fc-engineered antibodies as a novel class of TB therapeutics able to harness the protective functions of neutrophils to achieve disease control.

## INTRODUCTION

*Mycobacterium tuberculosis* (*Mtb*), the causative agent of tuberculosis (TB), remains the leading cause of death from a single bacterial infection globally, causing an estimated 1.5 million deaths in 2020^1^. As a result, novel therapeutic and vaccination strategies are urgently needed to slow the TB epidemic. To date, the majority of efforts to manipulate the immune response to drive protection or control of TB have focused on potentiating cell-mediated immunity, as CD4 T cells play a critical role in controlling TB^2–4^. Conversely, little work has focused on harnessing the diverse effector mechanisms of humoral immunity to combat TB.

A growing body of evidence supports a functional role for antibodies in TB control. Importantly, mice lacking B cells, the cellular source of antibodies, exhibit enhanced susceptibility to *Mtb* disease^5^. Moreover, several monoclonal and polyclonal antibody passive transfer studies have demonstrated that antibodies alone can limit TB disease and spread^6–12^. Of note, antibodies binding surface-exposed *Mtb* antigens have shown particular promise, demonstrating both an ability to reduce *Mtb* bacterial burden and prolong survival in treated animals^6,7,12–14^. More specifically, several studies have found that antibodies specific to capsular polysaccharides, the outermost portion of the bacteria, promote *Mtb* uptake and significant *Mtb* control *in vitro* and *in vivo*^6,7,12–14^. These data highlight the potential for exploiting antibodies that recognize abundant surface-exposed *Mtb* glycans to prevent TB disease.

While it is plausible that simple antibody blockade of surface-exposed antigens represents a mechanism of antibody action against *Mtb*, antibodies may also prompt bacterial clearance following surface opsonization via Fc-receptor engagement on local immune cells, allowing the activation of a diverse array of antimicrobial functions. Consistent with this model, antibody signaling via Fcγ receptors is required for optimal *Mtb* control *in vivo*^15^, strongly implicating the IgG Fc in protective immunity against TB. Moreover, antibody Fc functional profiles differ across individuals who control *Mtb* infection and those with active, uncontrolled infection^16^. However, the primary mechanism(s) exploited by the antibody Fc to promote *Mtb* control have not been thoroughly assessed.

In the present study, we sought to develop a more detailed understanding of the innate immune mechanism(s) that capsule-binding antibodies selectively leverage to restrict *Mtb*. Thus, we rationally engineered a library of antibody Fc-variants specific to α-glucan, an abundant, surface-exposed polysaccharide present in the *Mtb* capsule^17^. Each of these Fc-variants was designed to augment or dampen select antibody effector functions based on data from the monoclonal therapeutics field. We demonstrate that IgG Fc-engineering can significantly enhance the ability of α-glucan-specific antibodies to drive *Mtb* restriction *in vitro*. Fc-engineered α-glucan antibodies promoted *Mtb* restriction in a neutrophil-dependent manner, via the delivery of neutrophil survival signals able to mediate extended antimicrobial functions, raising the possibility of exploiting Fc-engineered antibodies as a novel class of therapeutics able to employ the antimicrobial activity of the innate immune system to drive TB control.

## RESULTS

### A wild-type IgG1 α-glucan antibody does not drive *Mtb* restriction *in vitro*

The capsule, the outer-most layer of *Mycobacterium tuberculosis*, is comprised primarily of glucan and arabinomannan, which represent approximately 80% and 20% of the capsular polysaccharides respectively^18,19^. While previous studies have demonstrated that glucan, the most abundant polysaccharide, elicits detectable antibody responses in mice and in humans^20–22^, it remains unclear whether antibodies to this highly abundant antigen possess antimicrobial activity. Thus, to begin to probe the antimicrobial function of α-glucan-specific humoral immunity, we exploited a monoclonal antibody, clone 24c5, previously shown to bind *Mtb*^20^.

Initially, the α-glucan-specific monoclonal antibody, 24c5, was generated as a human IgG1 monoclonal. Antigen binding to α-glucan was confirmed by ELISA and compared to that of an isotype control antibody (**Figure 1A**). Next, we sought to determine whether the α-glucan-specific IgG1 antibody was able to drive *Mtb* restriction *in vitro*. Human monocyte-derived macrophages (MDMs) were infected with a live/dead reporter strain of *Mtb* (*Mtb-live/dead*)^23^, followed by the addition of antibody to the *Mtb*-infected cells. After 96 hours, equivalent levels of intracellular *Mtb* killing was observed in wells containing the α-glucan-specific IgG1 antibody and the isotype control (**Figure 1B**). However, given that this assay interrogates the ability of antibodies to restrict bacterial growth solely in the presence of a previously infected macrophage, we next queried whether the antibody could restrict infection in human whole-blood – a system which captures the impact of multiple immune cell types and antibody functions at the time of bacterial exposure. Antibodies were added to fresh whole-blood from healthy human donors at the same time as an auto-luminescent *Mtb* reporter strain (*Mtb-276*)^24^. Luminescence readings were then taken every 24 hours over the course of 5 days to examine differences in *Mtb* growth curves across the α-glucan-specific and isotype control antibody conditions. Again, the α-glucan-specific IgG1 antibody did not exhibit any evidence of *Mtb* restriction (**Figure 1C**). Together, these data suggest that the α-glucan-specific IgG1 antibody did not mediate *Mtb* control *in vitro*.

**Figure 1:**
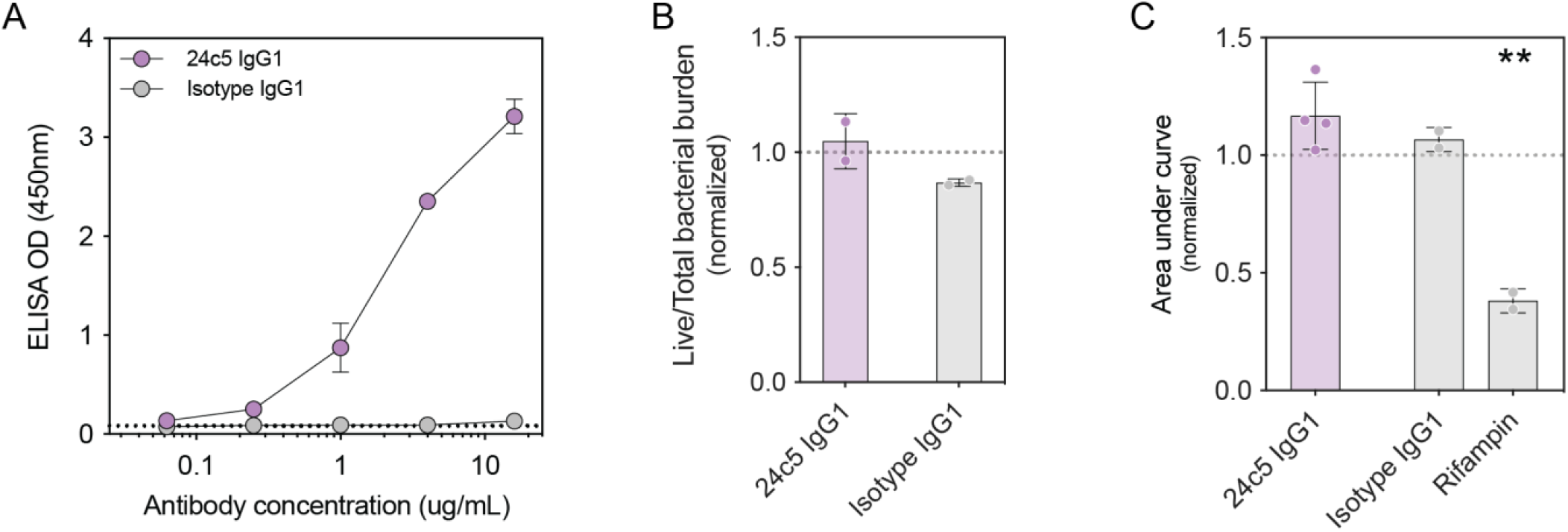
Wild-type IgG1 α-glucan antibody does not drive *Mtb* restriction *in vitro*. **(A)** Glucan (bovine liver glycogen) antigen-binding ELISA of the α-glucan-specific antibody clone, 24c5. **(B)** Macrophage *Mtb* restriction assay. Y-axis shows live (GFP) / total (mCherry) *Mtb* burden in human monocyte-derived macrophages normalized by the no antibody condition for the respective donor. Each point is the triplicate average from 1 human macrophage donor. Two-tailed, unpaired t test. Error bars show mean with standard deviation. **(C)** Whole-blood *Mtb* restriction assay. Y-axis is the area under the *Mtb-276* growth curve value normalized by the no antibody condition from the respective donor. Each point represents a triplicate average from one donor. One-way ANOVA with Dunnett’s correction comparing each antibody or antibiotic with the isotype IgG1 control antibody. Adjusted p-value: < 0.05 (*), < 0.01 (**), < 0.001 (***), < 0.0001 (****). Error bars show mean with standard deviation.

### Fc-engineered α-glucan antibodies display a range of functional activity

Functional and Fc-receptor binding differences related to differences in antibody Fc-glycosylation, were previously linked to enhanced antibody-mediated restriction of *Mtb* by IgG antibodies^16^. However, point mutations introduced into the antibody Fc domain at hotspots of Fc-receptor and complement protein binding provide a tightly regulated protein engineering framework that allows finer analysis of the relationship between select antibody Fc-mediated functions and microbial control^25–28^. Thus, to interrogate the impact of antibody functional Fc-profiles on the restrictive capacity of *Mtb*-specific antibodies, we developed a library of 52 Fc-engineered IgG antibody Fc-variants with identical antigen-binding fragments (Fabs) as the original 24c5 antibody clone using a high-throughput golden gate cloning approach (**Table 1**)^28^. Fc-variants included Fc-modifications known to modulate specific antibody functions such as antibody-dependent cellular cytotoxicity (ADCC), antibody-dependent phagocytosis by monocytes (ADCP), antibody-dependent complement deposition (ADCD), and serum half-life extension (**Table 1**)^28^. The 52 Fc-variants of 24c5 were produced and tested for their ability to bind to α-glucan by ELISA (**Figure S1A**). Each Fc-engineered antibody maintained binding comparable with that of the wild-type IgG1 antibody (**Figure S1A**), indicating that as expected, antibody Fc-modifications did not impede α-glucan binding activity.

**Table 1:**
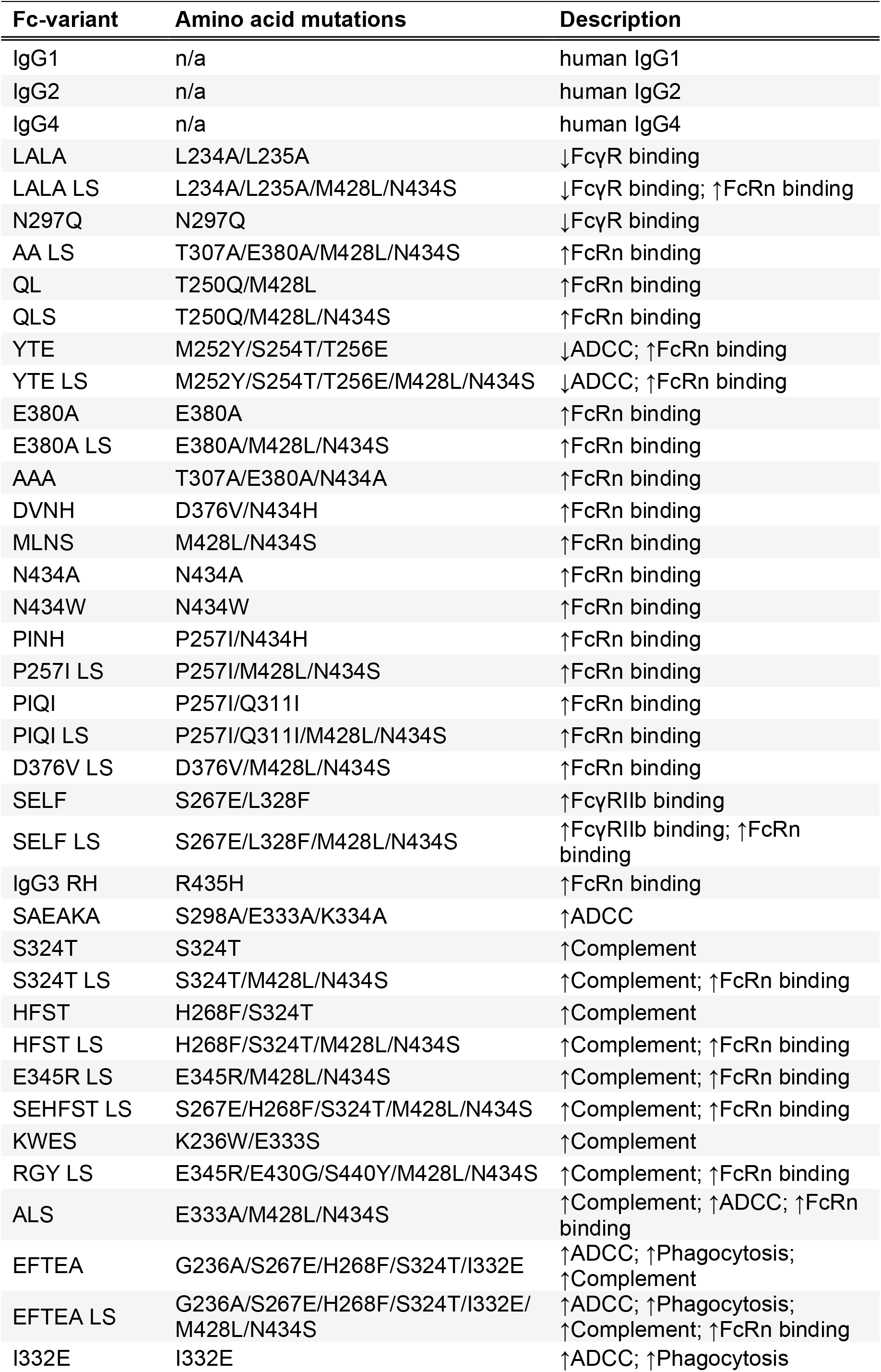
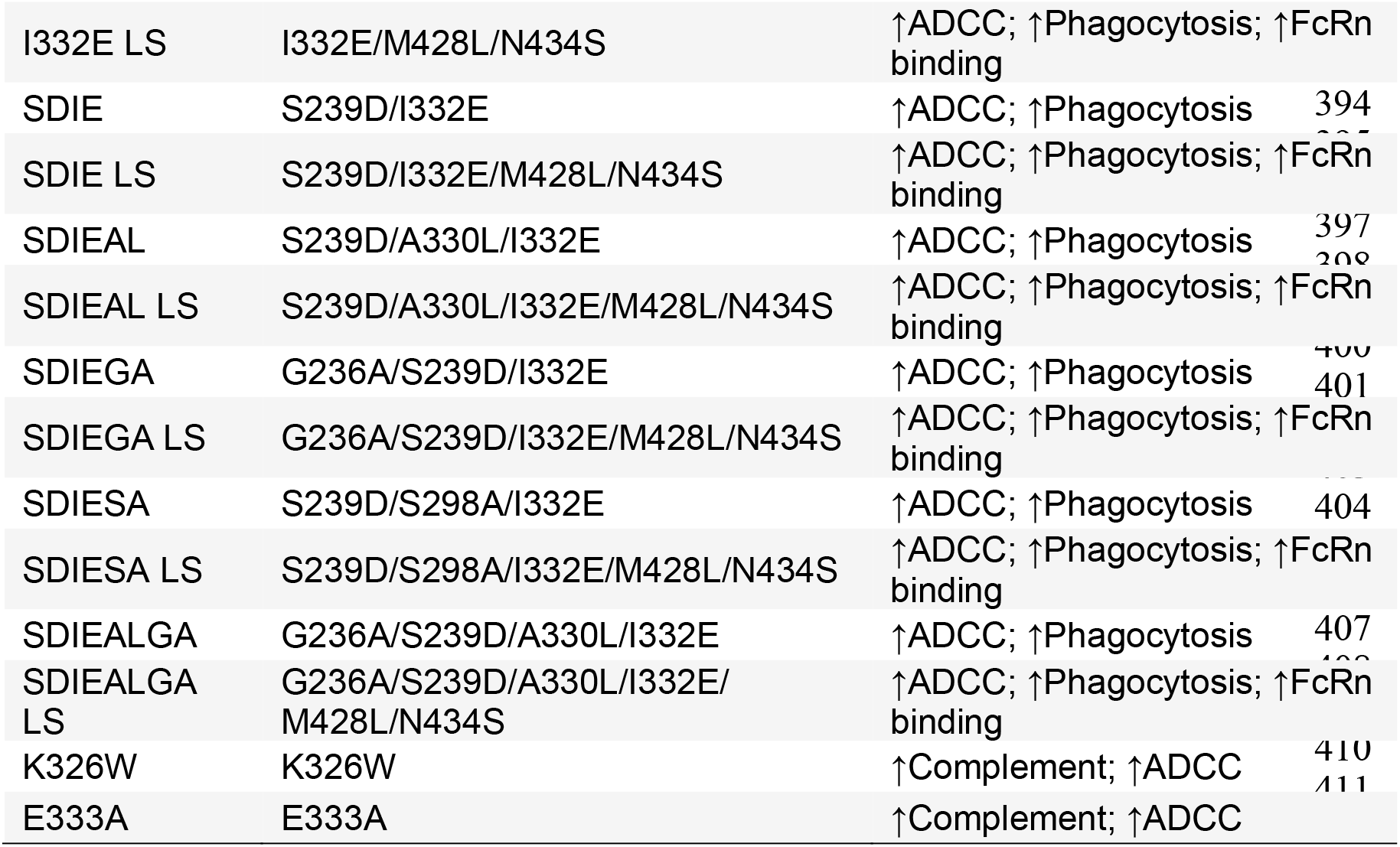
Library of Fc-engineered α-glucan antibodies. Amino acid mutations in the Fc domain are shown with a description of the expected functionality and the corresponding reference(s). Up arrow indicates increased functionality; down-arrow indicates decreased functionality. Antibody-dependent cellular cytotoxicity (ADCC), antibody-dependent cellular phagocytosis by monocytes (Phagocytosis), antibody-dependent complement deposition (Complement), binding to the neonatal Fc-receptor (FcRn binding), and binding to Fcγ receptors (Fcγ binding).

We next characterized the differential impact of each 24c5 Fc-variant on innate immune Fc-effector functions. Specifically, we probed the ability of each Fc-engineered antibody to drive NK cell activation, complement deposition, monocyte phagocytosis, and neutrophil phagocytosis in the presence of *Mtb* whole-cell lysate (**Figure 2** and **S1B**). As expected, in the antibody-dependent NK cell degranulation assay – a surrogate for ADCC – α-glucan Fc-variants engineered to have potent ADCC activity such as GASDELIE, SAEAKA, and I332E (**Table 1**)^28^, elicited increased NK cell degranulation (CD107a) and NK cell activation (IFNγ and MIP-1β secretion) compared to the wild-type IgG1 antibody (**Figure 2** and **S1B**). α-glucan Fc-variants such as KWES, K326W, and HFST, designed to have increased complement activity (**Table 1**)^28^, mediated increased complement component 3 (C3b) deposition compared to the wild-type IgG1 antibody (**Figure 2** and **S1B**). Similarly, α-glucan Fc-variants engineered to facilitate enhanced monocyte phagocytosis such as SDIEGA, SDIESA, and SDIEALGA

**Figure 2:**
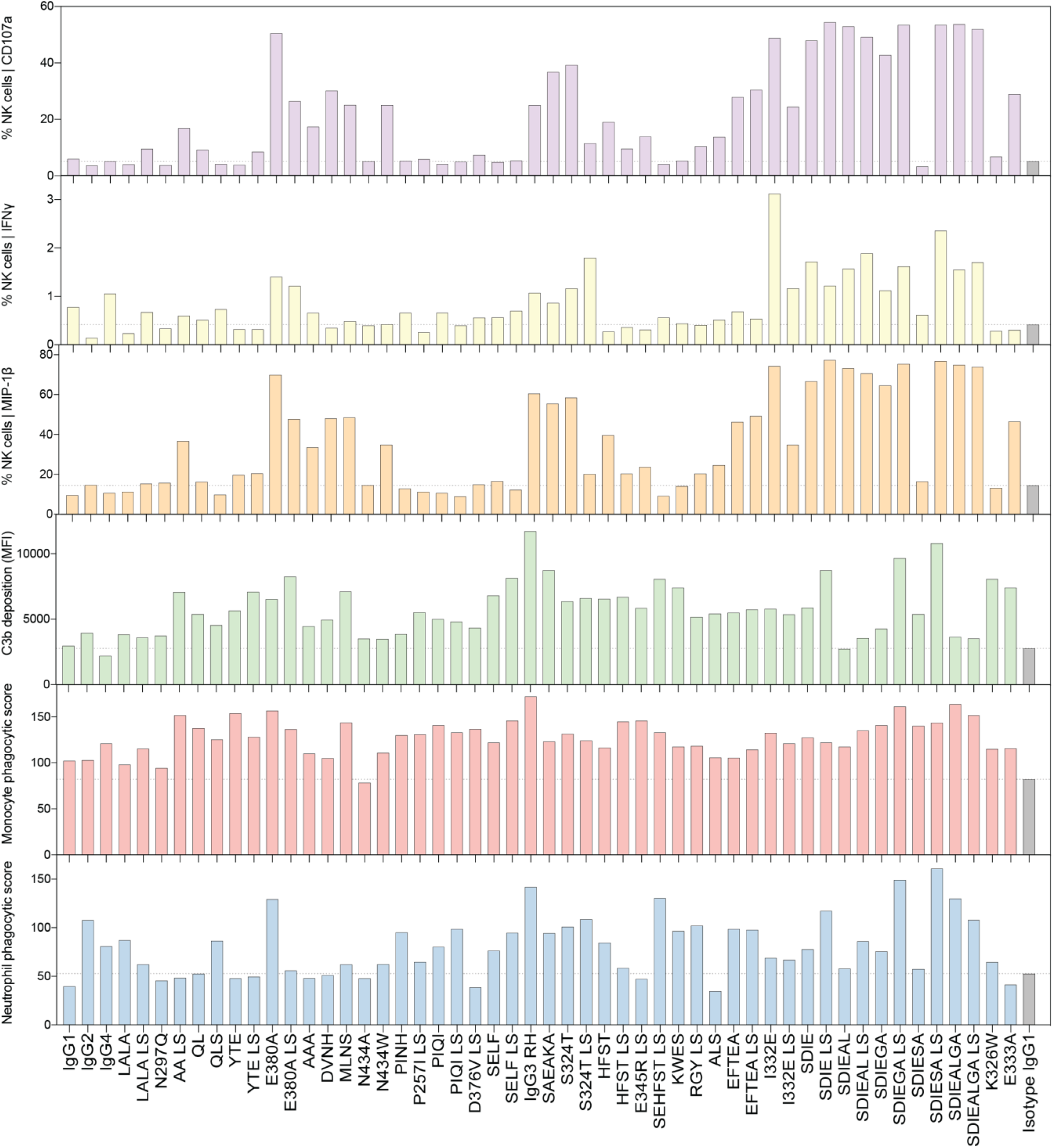
Fc-engineered α-glucan antibodies display a range in functional activity. The ability of each α-glucan Fc-variant to mediate (from top to bottom) NK cell degranulation (CD107a), NK cell secretion of IFNγ, NK cell secretion of MIP-1β, complement component 3b (C3b) deposition, phagocytosis by THP-1 monocytes, and phagocytosis by primary human neutrophils was experimentally determined. Each antibody was run in duplicate. Grey dotted line indicates the performance of an IgG1 isotype control antibody.

(**Table 1**)^28^, mediated increased monocytic uptake of *Mtb* whole-cell lysate-coated beads compared to the wild-type IgG1 antibody (**Figure 2** and **S1B**). Finally, while the ability of these Fc-variants to mediate neutrophil phagocytosis has not been as thoroughly assessed, several Fc-engineered α-glucan antibodies such as SEHFST LS, SDIESA LS, and IgG3 RH, drove robust neutrophil phagocytosis compared to the wild-type IgG1 antibody (**Figure 2** and **S1B**). From a combinatorial perspective, Fc-variants emerged with different combinations of antibody effector profiles (**Figure 2** and **S2B**). Of note, the N297Q variant, a non-glycosylated Fc-variant designed to have minimal affinity for Fcγ receptors^29,30^, exhibited limited activity across the functional profiling assays as expected (**Figure 2** and **S1B**). Collectively, these data indicate that Fc-engineering significantly shifts the functional profile of α-glucan-specific antibodies, driving the variable enhancement or diminution of several innate effector mechanisms of action, and providing a wide array of combinatorial functional responses to interrogate Fc-mediated restriction of *Mtb*.

### Fc-engineered α-glucan antibodies were down-selected by functional profile

To test the antimicrobial properties of α-glucan-specific antibodies with different Fc-effector profiles, we sought to reduce the number of α-glucan antibodies to a smaller set of variants that nonetheless maintained the functional heterogeneity present in the monoclonal library. To this end, the α-glucan Fc-variants were hierarchically clustered using all the functional profiling data. At least one variant was selected from each of the eleven clusters that emerged (**Figure 3A**), capturing the diversity in antibody functional profiles across the Fc-variant library. The ultimate down-selected 24c5 panel included Fc-variants with several types of functional activity (**Figure 3B**). For instance, the down-selected panel included YTE (which solely possessed monocyte phagocytic function), IgG3 RH (which displayed robust complement and phagocytic functions), I332E (which had potent NK activating properties), and N297Q (a largely non-functional and non-glycosylated Fc-variant) (**Figure 3B**). Thus, while the down-selected α-glucan Fc-variant panel is comprised of fewer variants, substantial heterogeneity was maintained, providing a robust starting point from which to define the relationship between antibody functional profiles and *Mtb* restriction.

**Figure 3:**
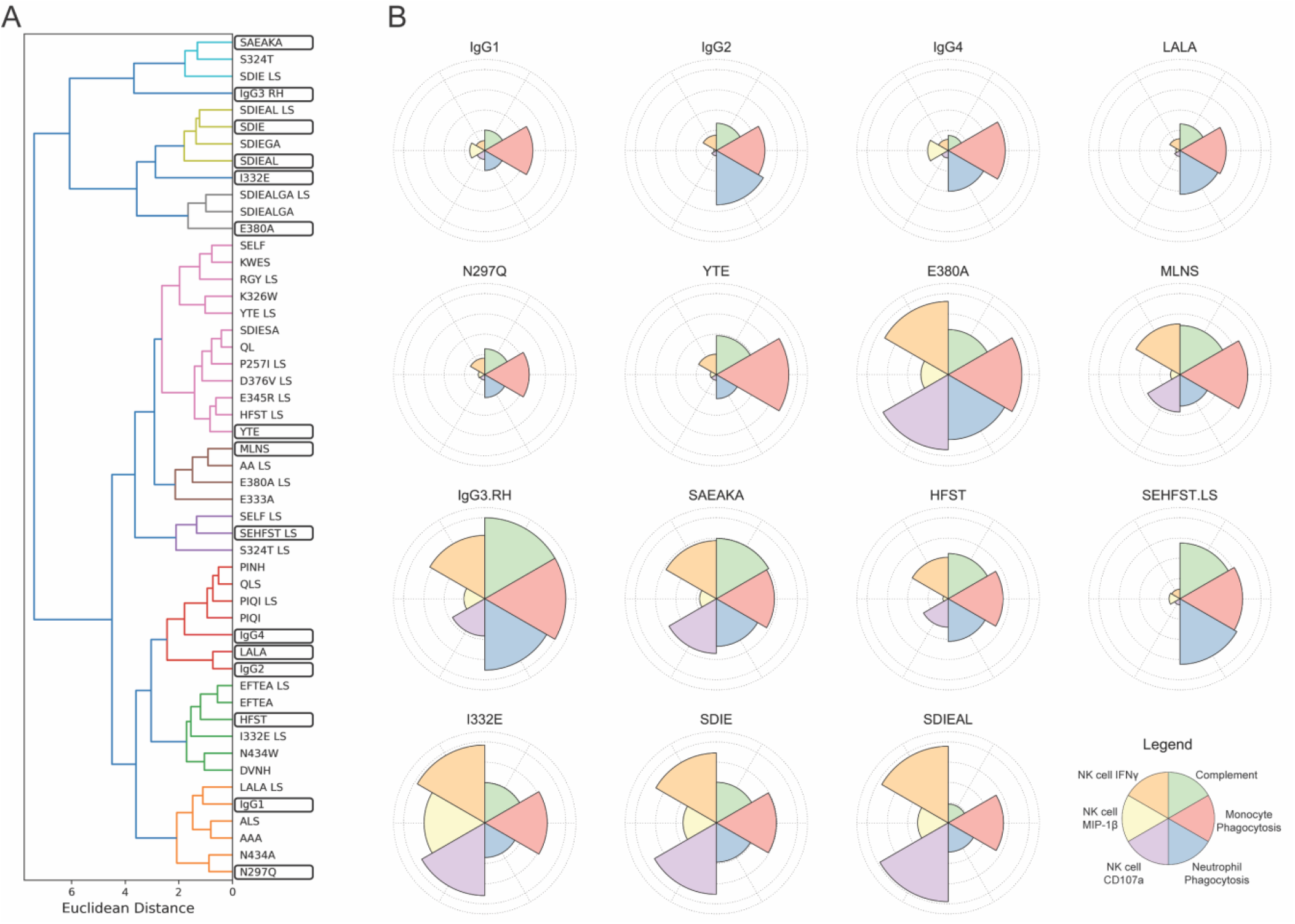
Hierarchical clustering utilized for the rational down-selection of Fc-engineered α-glucan antibodies. **(A)** Cluster dendrogram of the α-glucan Fc-variant panel following complete-linkage hierarchical clustering of the functional profiling data. At least one variant from each cluster (ellipsed) was selected for antimicrobial profiling. **(B)** Polar plots highlighting the functional profile of each α-glucan Fc-variant in the down-selected panel. Each pie piece indicates the magnitude of functionality in the respective assay relative to the entire panel. The α-glucan Fc-variants were max-scaled prior to polar plot visualization.

### Several Fc-engineered α-glucan antibodies drive *Mtb* restriction in whole-blood

While the wild type 24c5 IgG1 antibody did not significantly restrict *Mtb* (**Figure 1**), we next aimed to determine whether the addition of certain Fc-functions to the 24c5 antibody clone could augment bacterial restriction *in vitro*. Fresh whole-blood from healthy human donors was simultaneously infected with *Mtb-276*^24^, and treated with each 24c5 antibody. Notably, while the wild-type IgG1 antibody did not mediate significant *Mtb* restriction, 6 of the 15 down-selected Fc-engineered antibodies tested drove significant *Mtb* restriction in whole-blood compared to the IgG1 isotype control antibody (**Figure 4A**). Conversely, none of the Fc-variants tested drove significant intracellular *Mtb* killing in macrophages alone (**Figure S2**), suggesting the mechanistic involvement of additional peripheral blood cell types in antibody mediated killing *in vitro*.

**Figure 4:**
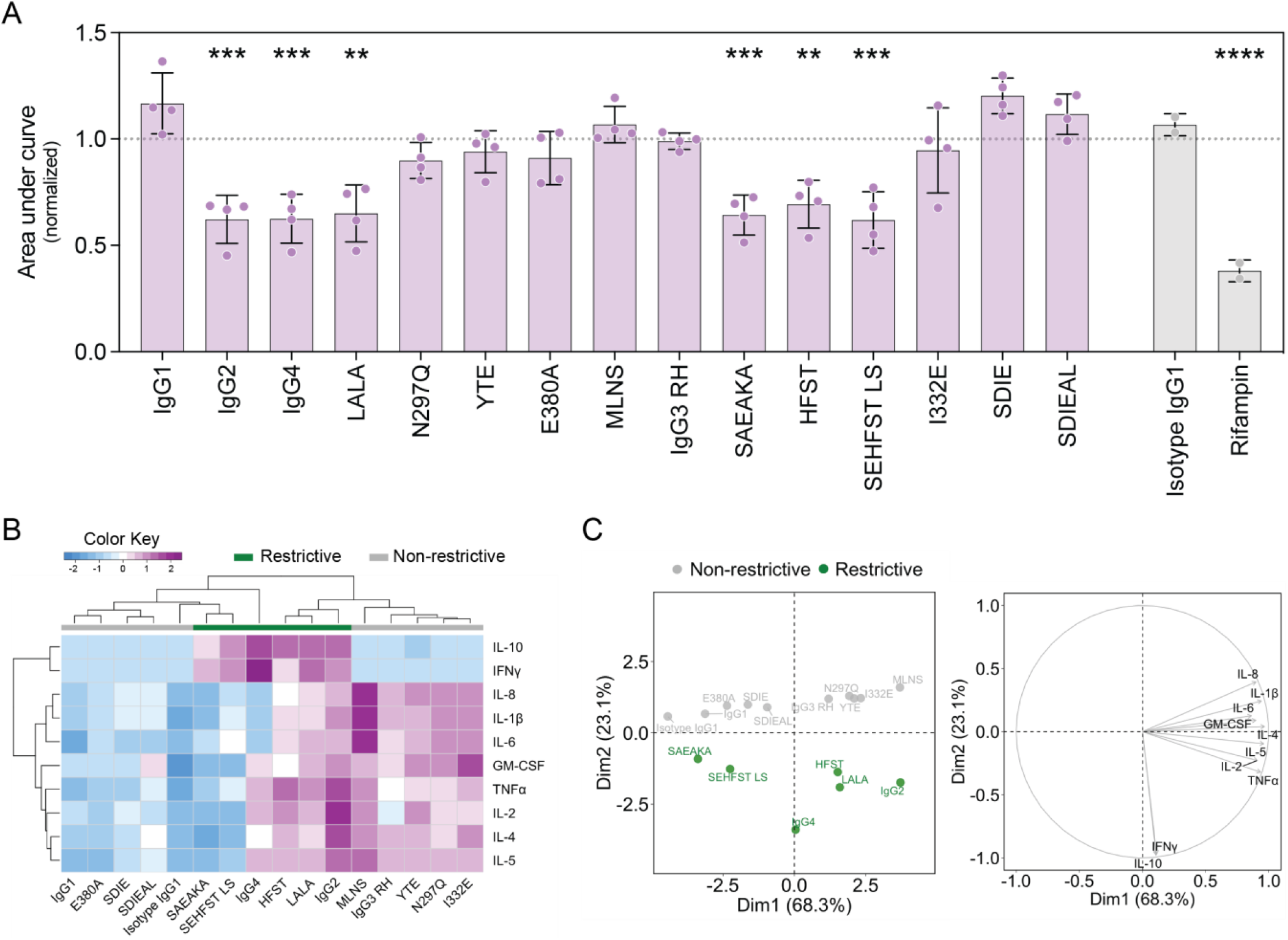
Several Fc-engineered α-glucan antibodies drive *Mtb* restriction in whole-blood. **(A)** Whole-blood *Mtb* restriction assay. X-axis shows the different α-glucan Fc-variants (25μg/mL), an IgG1 isotype control antibody as a negative control (25μg/mL), and the antibiotic rifampin as a positive control (0.25μg/mL). The y-axis is the area under the *Mtb-276* growth curve value normalized by the no antibody condition from the respective donor. Each point represents a triplicate average from one donor. **(B** and **C)** Cytokine Luminex using the whole-blood assay supernatant collected at 120 hours. Triplicate average from Donor A shown. One-way ANOVA with Dunnett’s correction comparing each antibody or antibiotic with the isotype IgG1 control antibody. Adjusted p-value: < 0.05 (*), < 0.01 (**), < 0.001 (***), < 0.0001 (****). Error bars show mean with standard deviation. **(B)** Clustered heatmap indicating the cytokine profile elicited by each α-glucan Fc-variant. Data were z-scored prior to heatmap visualization. **(C)** Principal component analysis of cytokine Luminex data. Left, score plot of the first two principal components. Right, loading plot of the first two principal components.

To explore the immune milieu driven by restrictive antibody treatment, we profiled cytokine levels 120 hours following *Mtb* infection and antibody treatment in whole-blood across multiple donors (**Figure 4B, C** and **S3**). Restrictive Fc-variants elicited a distinct cytokine profile, marked by a selective enrichment in secreted interferon-γ (IFNγ) and interleukin-10 (IL-10) compared to Fc-variants that were non-restrictive (**Figure 4B, C** and **S3**). These cytokine differences were significant at a univariate level (**Figure S3**). While there was heterogeneity in the secretion of additional cytokines including IL-6, IL-8, and IL-1β, these cytokines did not distinguish restrictive versus non-restrictive α-glucan antibody Fc-variants (**Figure 4C** and **S3**). Together, these data indicate that Fc-enhancement can result in *Mtb* restriction and highlight distinct cytokine/inflammatory cascades activated by restrictive antibodies.

### Fc-optimized α-glucan antibodies drive *Mtb* restriction in a neutrophil-dependent manner

To probe the mechanism(s) exploited by Fc-variants to restrict *Mtb* growth in whole blood, we next assessed the relationship between the whole-blood restriction assay and each of the functional profiling assays (**Figure 5A**). Notably, while several antibody effector functions positively correlated with one another, antibody-dependent neutrophil phagocytosis was the only Fc-effector function exhibiting a significant negative correlation with *Mtb* growth in whole-blood (**Figure 5A**). While we had anticipated that antibodies able to drive potent functions across multiple immune effector cell types would optimally restrict *Mtb*, unexpectedly, antibodies which were not broadly functional, and instead selectively induced neutrophil phagocytosis significantly restricted growth in whole-blood (**Figure 5B**). Conversely, antibody-dependent NK cell degranulation (CD107a) and activation (IFNγ and MIP-1β secretion) positively correlated with *Mtb* growth in whole-blood (**Figure 5A**).

**Figure 5:**
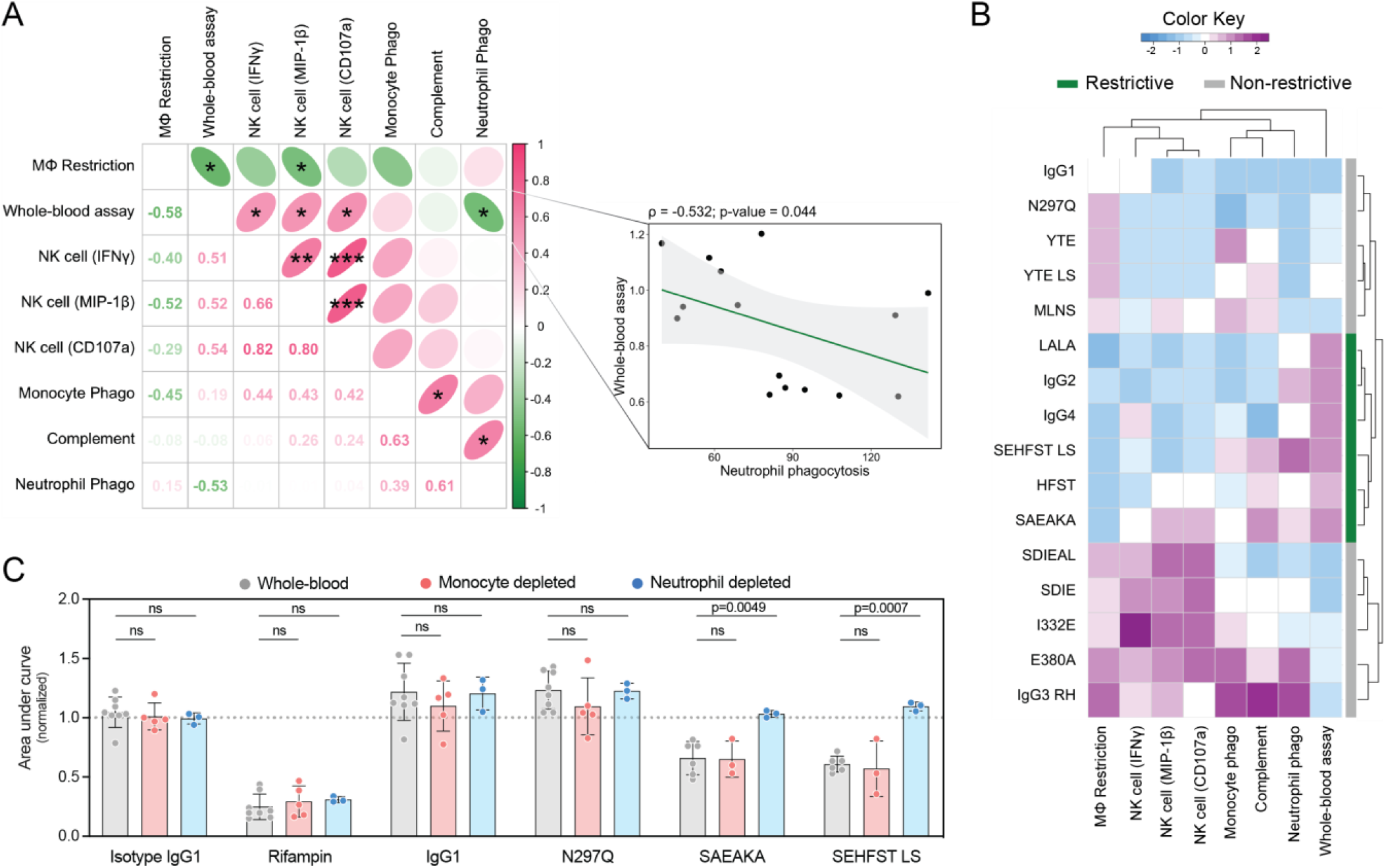
Fc-engineered α-glucan antibodies drive *Mtb* restriction in a neutrophil-dependent manner. **(A)** Spearman correlation matrix of the different functional and antimicrobial assays. Lower triangle indicates the spearman correlation coefficient for each relationship. Upper triangle contains ellipses which have their eccentricity parametrically scaled to the strength of the correlation for each relationship. Pairwise correlation between the normalized area under the curve in the whole-blood assay and the phagocytic score from the neutrophil phagocytosis assay is highlighted. **(B)** Clustered heatmap indicating the performance of each α-glucan Fc-variant in the down-selected panel across different functional and antimicrobial assays. Data were z-scored prior to heatmap visualization. **(C)** Whole-blood *Mtb* restriction assay with immune cell depletions. X-axis shows selected α-glucan Fc-variants (25μg/mL), an IgG1 isotype control antibody as a negative control (25μg/mL), and the antibiotic rifampin as a positive control (0.25μg/mL). Each antibody or antibiotic treatment was tested in whole-blood (grey), monocyte depleted blood (pink), or neutrophil depleted blood (blue). Y-axis is the area under the *Mtb-276* growth curve value normalized by the no antibody condition from the respective donor. Each point represents a triplicate average from one donor. One-way ANOVA with Dunnett’s correction comparing restriction in the monocyte and neutrophil depleted blood condition, with the whole blood condition for each treatment. Differences between blood conditions in the negative (Isotype IgG1) and positive (Rifampin) controls were not significant and thus are not shown. Error bars indicate mean with standard deviation.

To test the hypothesis that the engagement of neutrophils by restrictive Fc-engineered α-glucan antibody variants was key to *Mtb* restriction, neutrophils were depleted from whole-blood using commercially available magnetic beads prior to infection and prior to the addition of select Fc-engineered α-glucan antibodies. Strikingly, the SEHFST LS and SAEAKA α-glucan Fc-variants previously found to drive *Mtb* restriction, no longer restricted *Mtb* in the absence of neutrophils (**Figure 5C**). These α-glucan Fc-variants maintained their restrictive activity when monocytes were depleted instead of neutrophils (**Figure 5C**). As expected, neither the wild-type IgG1 nor the N297Q antibody variants drove *Mtb* restriction irrespective of neutrophil or monocyte immune cell depletion (**Figure 5C**). These data indicate that Fc-enhanced α-glucan antibodies leverage neutrophil rather than monocyte function to drive *Mtb* control.

### Fc-optimized antibody drives an upregulation of antimicrobial gene programs in neutrophils

Neutrophils have been associated with both protection and pathology in *Mtb* infection^31^. Neutrophils in some studies have been described as an early microbial reservoir in naïve hosts, controlling intracellular *Mtb* less effectively compared to macrophages^32,33^, and over the course of infection, neutrophil recruitment is associated with necrosis and caseation^31,34^. However, in other environments neutrophils have been found to have potent ability to clear *Mtb*, activity that is suppressed by granulocyte necrosis^35^. The observation that select Fc-engineered antibodies promoted *Mtb* restriction in a neutrophil-dependent manner, suggested that the presence of Fc-optimized antibodies at the time of bacterial exposure may trigger particular molecular circuits in neutrophils that direct neutrophil antimicrobial functions. Hence, we next performed single-cell RNA sequencing (scRNA-seq) to define the effects of Fc-optimized antibody treatment. Fresh whole-blood from three healthy human donors was infected with *Mtb* and treated with either the 24c5 IgG1 or the “restrictive” 24c5 SEHFST LS antibody variant. Conditions without monoclonal antibody (no Ab) and without *Mtb* (uninfected) were additionally included as controls. 24 hours following infection, we performed scRNA-seq analysis of the whole-blood cell populations under the various conditions.

Cells were clustered into one of twenty different cell subsets by their gene expression and visualized in low dimensional space through uniform manifold approximation and projection (UMAP) (**Figure 6A**). Following quality control, 7334 total cells and 906 neutrophils were recovered. Notably, the 24c5 SEHFST LS variant exhibited an increased proportional abundance of neutrophils compared to the other treatment conditions (**Figure 6B**). These data suggest increased survival of neutrophils in the presence of the Fc-optimized antibody as compared to the control conditions.

**Figure 6:**
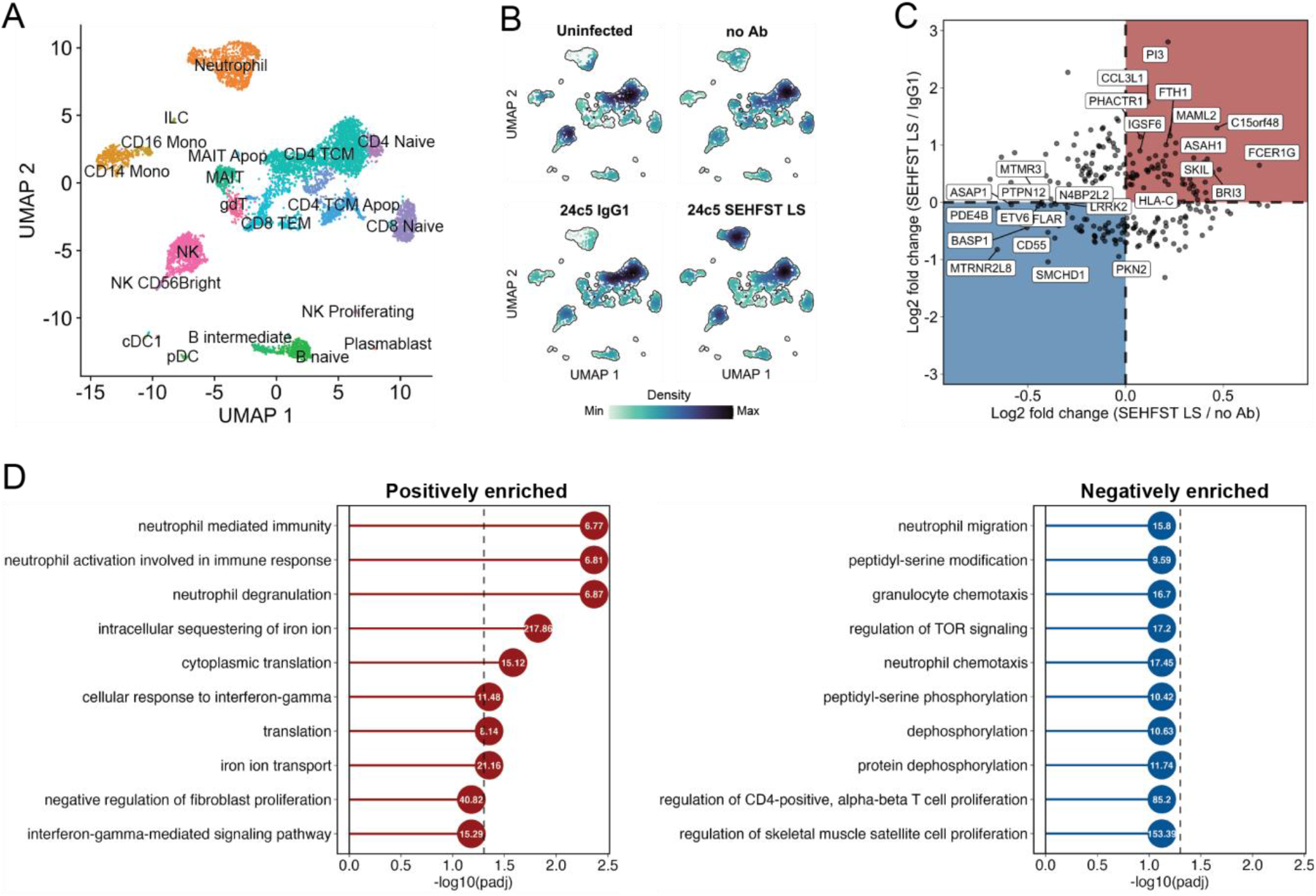
24c5 SEHFST LS drives the upregulation of antimicrobial gene programs in neutrophils. **(A)** UMAP visualization of all cells and cell subsets recovered following scRNA-seq. **(B)** UMAP depicting fractional abundance (density) in different cell types across treatment groups. **(C)** Neutrophil differential expression analysis. Genes consistently increased in the 24c5 SEHFST LS condition (red quadrant): (i) Mann-Whitney p-value < 0.1 and a log2 fold change > 0.25 compared to either the 24c5 IgG1 or no Ab condition, (ii) detected in a minimum fraction of 0.1 cells in either of the two conditions, and (iii) a log2 fold change > 0 compared to both the 24c5 IgG1 and no Ab conditions. Genes consistently decreased in the 24c5 SEHFST LS condition (blue quadrant): (i) Mann-Whitney p-value < 0.1 and a log2 fold change < -0.25 compared to either the 24c5 IgG1 or no Ab conditions, (ii) detected in a minimum fraction of 0.1 cells in either of the two conditions, and (iii) a log2 fold change < 0 compared to both the 24c5 IgG1 and no Ab conditions. **(D)** Gene list enrichment analysis using GO Biological Process gene sets. Left, GO terms enriched in red quadrant genes from panel C. Right, GO terms enriched in blue quadrant genes from panel C. Vertical dashed line indicates adjusted p-value of 0.05. Numbers on each circle show the odds ratio. Top ten GO terms by adjusted p-value shown.

Consistent with this, expression of Trappin-2 (*PI3*), an inhibitor of neutrophil elastase that may be associated with prevention of NETosis and alternative antimicrobial functions^36,37^, was increased in the 24c5 SEHFST LS condition compared to the 24c5 IgG1 and no Ab conditions (**Figure 6C**). Expression of a subunit of ferritin (*FTH1*), the major intracellular iron storage protein in eukaryotes^38^, was also higher in the setting of 24c5 SEHFST LS antibody treatment (**Figure 6C**), suggesting that the Fc-optimized antibody may modulate intracellular iron availability for the bacterium. Conversely, gene expression of *BASP1* (brain abundant signal protein 1), which is associated with cell death and senescence^39–41^, was decreased following 24c5 SEHFST LS antibody treatment (**Figure 6C**). Similarly, *CD55*, a well-characterized inhibitor of the complement pathway^42,43^, was consistently decreased in expression in the 24c5 SEHFST LS condition compared to the 24c5 IgG1 and no Ab conditions (**Figure 6C**), potentially pointing to a role for complement in antibody-mediated antimicrobial activity.

Gene Ontology (GO) analysis of the differentially expressed genes following 24c5 SEHFST LS antibody treatment (**Figure 6D**) found key antimicrobial circuits leveraged by neutrophils to drive control of intracellular pathogens including neutrophil degranulation^44,45^, sequestration of iron^46^, and the response to IFNγ^44^, to be significantly upregulated following 24c5 SEHFST LS treatment (**Figure 6D**). By contrast, analysis of the differentially expressed genes in the CD14+ monocytes from the 10X dataset revealed downregulation of the gene encoding toll-like receptor 2 (*TLR2*), and the GO biological processes involved in the response to cytokines following 24c5 SEHFST LS antibody treatment (**Figure S4**). Taken together, these data suggest that 24c5 SEHFST LS may divert the antimicrobial recognition and response to *Mtb* from the CD14+ monocyte to the neutrophil compartment, promoting neutrophil survival and the selective and sustained upregulation of antimicrobial gene programs that likely contribute to intracellular *Mtb* control.

## DISCUSSION

Here we utilized an Fc-engineering approach to, (i) determine whether rational Fc modification could enhance antibody restrictive capacity, and (ii) define the innate immune mechanism(s) that antibodies may leverage to restrict *Mtb*. We demonstrated that IgG Fc-engineering can significantly augment the ability of α-glucan antibodies to drive *Mtb* restriction *in vitro*, pointing to a novel strategy for the development of therapeutics to combat TB. Unexpectedly, Fc-engineered α-glucan antibodies drove *Mtb* restriction in a neutrophil-dependent manner. These data further suggest that antibodies can harness the antimicrobial potential of neutrophils to promote *Mtb* restriction by selectively rewiring neutrophils at the transcriptional level to shift from NETosis to alternative antimicrobial programs.

Activation of neutrophils represents a delicate and context-dependent balancing act. The robust destructive and inflammatory functions of neutrophils have been associated with the development of more severe TB disease late in infection^47,48^. Given that poorly functional antibodies are abundant late during active pulmonary *Mtb* infection^16,49^, it is plausible that these antibodies may exacerbate suboptimal neutrophil functions late in *Mtb* infection, rather than ameliorate TB disease. However, upon initial *Mtb* exposure, neutrophils are first recruited into the airways and lung-tissue at a time when adaptive immune responses are yet to develop, and thus in the presence of limited antigen-specific IgG antibodies. Conversely, in the present study, we observed that the pre-existence of *Mtb*-specific neutrophil-activating antibodies at the time of *Mtb* infection rewires neutrophils, promoting neutrophil survival and bacterial restriction. Our results show that the efficacy of neutrophils against mycobacteria is highly dependent on whether the interaction involves immunoglobulin opsonins and the type of engagement of Fc receptors. Consequently, the efficacy of neutrophils in the control of mycobacterial infection depends on the quality and quantity of specific antibodies available. Since the humoral response to mycobacterial antigens is complex and dependent on the duration of infection, it is possible that the efficacy of neutrophils against mycobacteria also depends on the kinetics of the humoral response and varies with time. Vaccines that elicit antibodies that facilitate neutrophil activity against mycobacteria could protect by enhancing the efficacy of these innate immune response cells, which are often first responders to sites of infection. In this respect, recent analyses of bronchoalveolar lavage fluid from non-human primates immunized with protective intravenous BCG identified a significant expansion of antibodies able to arm neutrophil activity in the lung^50^.

Given the limited tools to study the physical milieu and dynamics of the *Mtb* phagosome, the primary replicative niche of the bacterium^51^, defining the mechanistic opportunities to develop next generation therapeutics to *Mtb* has been challenging. Nevertheless, the combination of Fc-engineering with single cell transcriptomics pointed to novel and unexpected Fc-mediated antibody therapeutic strategies as alternatives in an expanding world of antibiotic resistance^1^. Specifically, transcriptomic analysis pointed to the need for neutrophil survival and degranulation for optimal antibody-mediated *Mtb* restriction. While macrophages and monocytes shunt phagosomal contents into the endocytic maturation pathway, gradually converting the phagosome into a phagolysosome, neutrophils instead possess a myriad of preformed granules containing antimicrobial peptides and lytic enzymes that rapidly fuse with the phagosome following Fcγ-mediated uptake^45^. Thus, it is plausible that neutrophil survival, if long enough to permit the unique phagosome-targeted degranulation process downstream of Fcγ-mediated phagocytosis in neutrophils, may contribute to intracellular *Mtb* control. Finally, more complex multicellular functions through antigen presentation and cytokine release may also promote control^44,52,53^. Indeed, the significant increase in IFNγ release induced by restrictive Fc-variants, may hint at the importance of the induction of an antimicrobial cytokine response cascade that may contribute to mycobacterial restriction.

Invulnerable to antibiotic resistance, antibody-based therapeutics against TB represent an appealing therapeutic modality in the age of increased drug resistant TB cases globally^1^. Thus, while little investigative effort has focused on harnessing humoral immunity to combat TB, our work demonstrates the value of exploiting Fc-engineering and single cell transcriptional analysis to identify novel antibody-mediated mechanisms of *Mtb* control and to inspire next generation antibody-based therapeutic design.

## ACKNOWLEDGEMENTS

Thanks to Eric Rubin and Jeffrey Wagner for sharing the luminescent *Mtb* reporter strain used in this study. Thanks to the Laboratory for Systems Pharmacology at Harvard Medical School for allowing the use of their automated microscope. This work was made possible with help from the Harvard University Center for AIDS Research (CFAR), an NIH funded program (P30 AI060354).

## Author contributions

Conceptualization: EBI, RL, PSG, SMF, GA. Methodology: EBI, JMP, RL, PSG, JS, JCH, SMF, GA. Software: EBI, JMP. Validation: EBI, AW, MS, SS. Formal Analysis: EBI, JMP. Investigation: EBI, JMP, PSG, JS, SS, WK, JCH, EvW. Resources: AW, MS, SS, WK, AC, LC. Data Curation: EBI, JMP, PSG. Writing Original Draft: EBI. Review and Editing: all authors. Visualization: EBI. Supervision: BB, LC, SMF, GA. Project Administration: SMF, GA. Funding Acquisition: EBI, BB, LC, SMF, GA.

## Data Availability

All data associated with this study are available at: XXXXXXX

## Code Availability

Scripts to reproduce the computational analyses presented in the manuscript are available at: XXXXXXX

## MATERIALS AND METHODS

### Fc-engineering

A golden gate cloning strategy was employed for REFORM antibody Fc-engineering as described previously^28,54^. In short, BsaI restriction sites flanking the sequences of the different antibody domains [variable heavy (VH), variable light (VL), constant heavy (which was distinct for each Fc-variant), constant light], as well as a furin 2A domain were inserted. BsaI generates a set of unique overhangs, allowing the antibody expressing plasmid to be generated in a single-step digestion/ligation reaction. The furin 2A site mediates self-cleavage of the polypeptide, and thus expression of the entire antibody from a single open reading frame^55^. Each plasmid was sequenced to confirm the accuracy of golden gate assembly.

### Antibody expression and purification

Antibody VL plasmids were co-transfected with each antibody VH plasmid at a 1:1 ratio into CHO cells. Antibody was purified from the cell supernatant using a Prosep-vA Ultra Protein A resin. Antibody was then dialyzed and concentrated in phosphate buffered saline (PBS). Concentration was determined by ELISA through comparison with a known standard.

### Glucan ELISA

ELISA plates (Thermo Fisher, NUNC MaxiSorp flat bottom) were coated with 50µL of bovine liver glycogen (Milipore Sigma, G0885) at 100µg/mL overnight at 4°C. The plates were washed with PBS and blocked with 5% bovine serum albumin (BSA)-PBS for 2 hours at room temperature (RT) on an orbital shaker. The plates were washed with PBS, then 80µL of antibody was added in 4-fold dilutions ranging from 16µg/mL to 0.0625µg/mL and the plate was incubated for 1.5 hours at RT on an orbital shaker. The plates were washed with PBS, then 80µL of secondary anti-human Igκ light chain HRP-conjugated antibody (ThermoFisher, A18853) diluted 1:1000 in 0.1% BSA-PBS was added. The plates were incubated for 1 hour at RT on an orbital shaker. The plates were washed with PBS, then 80µL per well of TMB (ThermoFisher, 34029) was added. The reaction was stopped using 2N H_2_SO_4_, and absorbance was measured at 450nm on a plate reader (Tecan Infinite M1000 Pro).

### Antibody-dependent cellular phagocytosis (ADCP)

ADCP was performed as described previously^50,56^ with slight modifications. 250µg of *Mtb* H37Rv whole-cell lysate (BEI, NR-14822) was first treated with sodium acetate (25µL NaOAc at 1M and pH 5.5) and sodium periodate (55µL NaIO_4_ at 50mM), and incubated for 90min at RT in the dark. Next, sodium bisulfate (30µL NaHSO_4_ at 0.8M and diluted in 0.1M NaOAc) was added to block the oxidation reaction for 5min at RT in the dark. The oxidized whole-cell lysate solution was transferred to a new tube, then 55µL of hydrazide biotin at 50mM (Sigma, 21339), 25µL of 1M NaOAc, and 175µL of ddH _2_O were then added to the oxidized whole-cell lysate, and incubated for 2hr at RT. Following the incubation, excess biotin was removed using Amicon Ultra 0.5L columns (3K, Millipore Sigma). The biotinylated whole-cell lysate was then added to FITC neutravidin beads (ThermoFisher, F8776) at a ratio of 5µg antigen:1µL beads, and incubated shaking overnight at 4°C. Whole-cell lysate-coated beads were centrifuged at 16,000 x g for 5 min, resuspended in 1mL 5% BSA-PBS, and incubated shaking at RT for 1hr to block. Whole-cell lysate-coated beads were then resuspended in 5% BSA-PBS such that the starting 1µL of beads were in 100µL of solution. A 10µL volume of whole-cell lysate-coated beads were then incubated with 40µL of each monoclonal antibody at 0.025µg/mL (1µg antibody total) for 2hr at 37 °C to form immune-complexes. After the immune-complexes were washed with PBS, THP-1 monocytes (5.0 × 10^4^ per well) were added and incubated with the immune-complexes for 16hr at 37 °C. Fluorescent bead uptake was measured in 4% paraformaldehyde (PFA) fixed cells by flow cytometry on a BD LSRII (BD Biosciences) and analyzed by FlowJo 10.3. Phagocytic scores were calculated as: ((%FITC positive cells) x (geometric mean fluorescence intensity of the FITC positive cells)) divided by 10,000. Samples were run in duplicate.

### Antibody-dependent neutrophil phagocytosis (ADNP)

ADNP was performed as described previously^50,57^, with minor modifications. *Mtb* H37Rv whole-cell lysate was oxidized, biotinylated, coupled to FITC neutravidin beads (ThermoFisher, F8776), incubated with antibody, and washed as described in the previous section for ADCP. Next, fresh blood collected from healthy donors in acid citrate dextrose anti-coagulant tubes was added at a 1:9 ratio to ACK lysis buffer (Quality Biological, 10128-802) and incubated for 5 minutes at RT. Leukocytes were washed with PBS and resuspended in R10 medium – RPMI (Sigma), 10% fetal bovine serum (Sigma), 10mM HEPES (Corning), 2mM L-glutamine (Corning) – at a concentration of 2.5 × 10^5^ cells/mL. 5 × 10^4^ leukocytes per well were added to the immune-complexed beads and incubated for 1hr at 37°C. The cells were stained with anti-human CD66b-Pacific Blue (BioLegend), washed with PBS, then fixed with 4% PFA. Bead uptake was measured by flow cytometry on a BD LSRII (BD Biosciences) and analyzed by FlowJo 10.3. Phagocytic scores were calculated in the CD66b positive cell population. Samples were run in duplicate.

### Antibody-dependent complement deposition (ADCD)

ADCD was performed as described previously^28,58^, with minor modifications. *Mtb* H37Rv whole-cell lysate was oxidized, biotinylated, coupled to red fluorescent neutravidin beads (ThermoFisher, F8775), incubated with antibody, and washed as described in the ADCP section. Next, guinea pig complement (Cedarlane, CL4051) diluted in magnesium and calcium containing veronal buffer (Boston Bioproducts) was added to the immune-complexed beads and incubated for 20min at 37°C. Beads were then washed in 15mM EDTA-PBS and stained with FITC-conjugated anti-guinea pig C3 antibody (MP Biomedicals, MP0855385). C3 deposition onto beads (FITC MFI of all beads) was evaluated by flow cytometry on a BD LSRII (BD Biosciences) and analyzed by FlowJo 10.3. Samples were run in duplicate.

### Antibody-dependent NK cell activation (ADNKA)

ADNKA was performed as described previously^50^, with minor modifications. ELISA plates (Thermo Fisher, NUNC MaxiSorp flat bottom) were coated with 250ng/well of bovine liver glycogen (Milipore Sigma, G0885) overnight at 4°C. The plates were washed with PBS and blocked with 5% BSA-PBS at RT for 2hr. The plates were washed with PBS, and 40µL of each monoclonal antibody at 0.025µg/mL (1µg antibody total) was added and incubated for 2hr at 37°C. One day prior to adding antibody, NK cells were isolated from healthy human donors using the RosetteSep human NK cell enrichment cocktail (Stemcell) and incubated overnight at 1.5 × 10^6^ cells/mL in R10 media with 1ng/mL of IL-15 (Stemcell) at 37°C. After the 2hr incubation on the day of the assay, the ELISA assay plates were washed with PBS, and 50,000 NK cells, 2.5uL PE-Cy5 anti-human CD107a (BD), 10uL GolgiStop (BD), and 0.4uL Brefeldin A (5mg/mL, Sigma) were added to each well. The ELISA assay plates were incubated for 5hr at 37°C. After the incubation, cells were stained for surface expression with Alexa Fluor 700 anti-human CD3, PE-Cy7 anti-human CD56, and APC-Cy7 anti-human CD16 (all from BD). The cells were washed with PBS, then fixed using Perm A and Perm B (Invitrogen). The Perm B solution contained PE anti-human MIP-1β and FITC anti-human IFNγ (both from BD) for intracellular cytokine staining. The cells were washed, and the fluorescence of each marker was measured on a BD LSR II flow cytometer (BD Biosciences) and analyzed by FlowJo 10.3. The assay was performed in biological duplicate with NK cells from two donors.

### Macrophage restriction assay

*In vitro* macrophage *Mtb* survival was measured as described previously^16,50^, with slight modifications. CD14 positive cells were isolated from healthy donors using the EasySep CD14 Selection Kit II (Stemcell). CD14 positive cells were matured for 7 days into human monocyte-derived macrophages (MDMs) in R10 media without phenol in low adherent flasks (Corning), then plated at 50,000 cells per well in glass bottom, 96-well plates (Greiner) 24 hours prior to infection. *Mtb* H37Rv with constitutive mCherry and anhydrotetracycline inducible green fluorescent protein (GFP) expression (*Mtb-live/dead*)^23^, was cultured in log phase and filtered through a 5μm filter (Milliplex) prior to MDM infection at a multiplicity of infection of 1 overnight at 37°C. Infected MDMs were washed with PBS, and 200µL of monoclonal antibody at 50µg/mL in R10 without phenol was added and incubated at 37°C. 3 days after infection, anhydrotetracycline (Sigma) was added to the infected MDMs at 200 ng/mL and incubated for 16hr at 37°C. Cells were then fixed with 4% PFA and stained with DAPI. Data were analyzed using the Columbus Image Data Storage and Analysis System. *Mtb* survival was calculated as the ratio of live to total bacteria within macrophages in each well. *Mtb* survival for each condition was normalized by *Mtb* survival in the no antibody condition. The assay was performed in technical triplicate and in MDMs from two donors.

### Whole-blood restriction assay (WBA)

Whole-blood from healthy human donors was collected the day of the experiment in acid citrate dextrose anti-coagulant tubes. An auto-luminescent H37Rv *Mtb* reporter strain (*Mtb-276*)^24^, was cultured in log phase, washed, then resuspended in R10 without phenol. Whole-blood was simultaneously infected with *Mtb-276* at 1.0×10^6^ bacteria per mL of blood, and treated with antibody in a white, flat-bottom 96-well plate (Greiner). Final concentration of antibody treatments was 25μg/mL. Final concentration of rifampin (Sigma) positive control was 0.25μg/mL. At each timepoint – immediately after infection, and every 24 hours post-infection until 120 hours – the samples in each well were mixed, and a luminescence reading was taken (Tecan Spark 10M) to generate *Mtb* growth curves in the presence of each treatment. *Mtb* restriction in whole-blood was calculated as the area under the curve for each condition. Area under the curve values were computed in GraphPad Prism (version 8.4.0). The assay was performed in technical triplicate and in blood from multiple donors.

### Blood immune cell depletions

StraightFrom Whole Blood CD66b (Miltenyi, 130-104-913), CD14 (Miltenyi, 130-090-879), or Basic unconjugated (Miltenyi, 130-048-001) microbeads were first buffer-exchanged into 2mM EDTA-PBS. Specifically, a Whole Blood Column (Miltenyi, 130-093-545) was attached to the MidiMacs (Miltenyi) and primed with 3mL of Separation Buffer (PBS with 0.5% BSA and 2mM EDTA). 2mL of microbeads was added to the primed Whole Blood Column and the flow-through was discarded. After washing the column 3 times with 500μL of PBS, the microbeads were eluted with a final volume of 1mL in 2mM EDTA-PBS. On the day of the assay, whole-blood from healthy human donors was collected in acid citrate dextrose anti-coagulant tubes. 100μL of microspheres was added per mL of blood, and the solution was incubated for 20min at 4°C. During the incubation, Whole Blood Columns were attached to the MidiMacs, and primed with 3mL of Separation Buffer followed by 3mL of R10 media. Following the incubation, the microsphere-treated blood was added to the column, and the flow-through was collected and used in the whole-blood restriction assay described in the previous section.

### Cytokine Luminex

At the final timepoint of the whole-blood restriction assay (120hr), the assay plates were centrifuged at 800 x g for 5min, then 110μL of supernatant was collected from each well and transferred into a separate 96-well plate which was stored at -20°C until further use. Supernatants were thawed and twice filtered using 0.2µm 96-well filter plates (Millipore Sigma, CLS3508) for removal from the biosafety level 3 laboratory space. The abundance of select cytokines was then measured in 50μL of the supernatants with a Human Cytokine Magnetic 10-Plex Panel (ThermoFisher, LHC0001M) according to the instructions of the manufacturer. Median fluorescence intensity (MFI) for each analyte was measured using a FlexMap 3D (Luminex Corp.). Samples were measured in technical triplicate in blood from 2 donors.

### Single-cell RNA-sequencing (scRNA-seq)

Whole-blood *Mtb* infection and the addition of antibody was performed as described above for the whole-blood restriction assay. Blood from three healthy donors was utilized. 24 hours following infection and the addition of antibody, blood from each donor was added at a 1:9 ratio to ACK lysis buffer (Quality Biological, 10128-802) and incubated for 10min at RT. Cells were centrifuged at 400g for 5min, and the supernatant was discarded. 10mL of ACK lysis buffer was added to the cell pellet to repeat the lysis procedure. Cells were centrifuged at 400g for 5min, and the supernatant was discarded. Samples were washed twice with PBS buffer and counted before MULTI-seq barcoding as described^59^. In brief, samples were barcoded with 2.5µM of the LMO anchor and barcode for 5min on ice in PBS before adding 2.5µM of the LMO co-anchor and incubating for an additional 5min. Samples were quenched with 1% BSA in PBS and washed once. Samples were pooled and 0.5U/µL RNase inhibitor (Roche) was added. Pooled samples were then loaded into two lanes using the 10X Genomics NextGEM Single Cell 3’ kit v3.1 per the manufacturer’s protocol. cDNA was inactivated at 95°C for 15min prior to biosafety level 3 removal for library construction. Libraries were sequenced on a NextSeq500 (Illumina). FASTQ files were processed using CellRanger v6.1.2 to generate gene expression count matrices and deMULTIplex to generate LMO barcode count matrices.

### scRNA-seq data analysis

LMO barcode and gene expression count matrices were analyzed using R (v4.0.3) and Seurat (v4.0.0). Cells were identified using emptyDrops (DropletUtils) were demuxed using HTODemux (Seurat) and hashedDrops (DropletUtils). Each lane was subject to demultiplexing and quality control separately then merged for downstream analyses. Cells with less than 300 unique genes detected were excluded. Additional cells were excluded based on the assessment of cluster-specific technical metrics (percent of mitochondrial reads per cell and number of UMIs per cell). Counts were normalized using the default parameters from NormalizeData (Seurat), i.e. scaling by 10,000 and log normalization. 3,000 variable features were used for PCA. SLM clustering was performed using FindClusters (Seurat) on the shared nearest neighbor graph generated from FindNeighbors (Seurat) using 30 principal components and k=20. Cell type annotation was based on expert annotation and predicted cell type labels from the PBMC dataset in Azimuth using FindTransferAnchors and TransferData (Seurat). Marker gene statistics were calculated using wilcoxauc (presto). Gene expression matrices are accessible on GEO (GSEXXXXXX). Genes consistently increased following 24c5 SEHFST LS antibody treatment were defined as those: (i) with a Mann-Whitney p-value < 0.1 and a log2 fold change greater than 0.25 compared to either the 24c5 IgG1 or no Ab condition, (ii) detected in a minimum fraction of 0.1 cells in either of the two conditions, and (iii) with a log2 fold change greater than 0 compared to both the 24c5 IgG1 and no Ab conditions. Genes consistently decreased following 24c5 SEHFST LS antibody treatment were defined as those: (i) with a Mann-Whitney p-value < 0.1 and a log2 fold change less than -0.25 compared to either the 24c5 IgG1 or no Ab conditions, (ii) detected in a minimum fraction of 0.1 cells in either of the two conditions, and (iii) a log2 fold change less than 0 compared to both the 24c5 IgG1 and no Ab conditions.

### Gene ontology analysis

Gene Ontology analysis was performed using the Enrichr web-based platform^60^. Input genes for the 24c5 SEHFST LS positive enrichment analysis included genes consistently increased following 24c5 SEHFST LS antibody treatment as compared to both the 24c5 IgG1 and the no Ab conditions (**Figure 6C**, red quadrant). Input genes for the 24c5 SEHFST LS negative enrichment analysis included genes consistently decreased following 24c5 SEHFST LS antibody treatment as compared to both the 24c5 IgG1 and the no Ab conditions (**Figure 6C**, blue quadrant). Gene Ontology Biological Process was the gene set source. Top 10 gene sets ranked by Benjamini-Hochberg adjusted p-value are shown^61^. Gene sets with an adjusted p-value < 0.05 were considered significant.

### Multivariate analyses

Complete-linkage hierarchical clustering was performed on the z-scored antibody functional data using the SciPy library in Python (version 3.8.8). Polar plots were generated on the max-normalized antibody functional data using the ggplot2 (version 3.3.5) package in R (version 4.1.1). Clustered heatmaps of the z-scored antibody functional data were generated using the gplots (version 3.1.1) package in R (version 4.1.1). Principal component analysis on the z-scored cytokine data was performed using the factoextra (version 1.0.7) and ggplot2 (version 3.3.5) packages in R (version 4.1.1). Spearman correlations analyses were performed using the corrplot (version 0.92) package in R (version 4.1.1).

### Statistics

For the macrophage restriction assay, whole-blood restriction assay, and cytokine Luminex assay, one-way ANOVA tests were implemented with Dunnett’s correction comparing each antibody with the isotype IgG1 control antibody. For the blood immune cell depletion assay, a one-way ANOVA with Dunnett’s correction was implemented, comparing restriction in the monocyte and neutrophil depleted blood condition, with the whole-blood condition for each antibody treatment. These statistics were performed in GraphPad Prism (version 8.4.0). Spearman correlations between antibody functional assays were computed in R (version 4.1.1).

