## Supplemental Materials for "Fc-engineered antibodies leverage neutrophils to drive control of *Mycobacterium tuberculosis*"

### SUPPLEMENTAL FIGURES

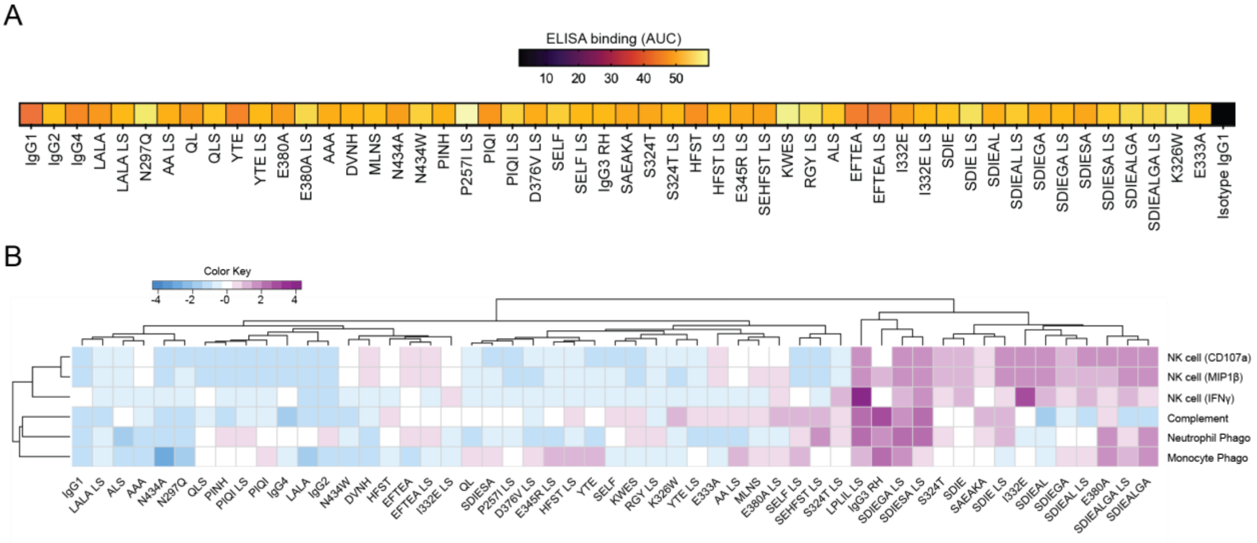


**Figure S1: Functional profiling of Fc-engineered α-glucan antibodies.** (**A**) Glucan (bovine liver glycogen) antigen-binding ELISA of the α-glucan-specific Fc-variant panel. Area under the dilution curve is plotted in heatmap. (**B**) Clustered heatmap indicating the performance of each α-glucan Fc-variant in the functional profiling assays. Data were z-scored prior to heatmap visualization.

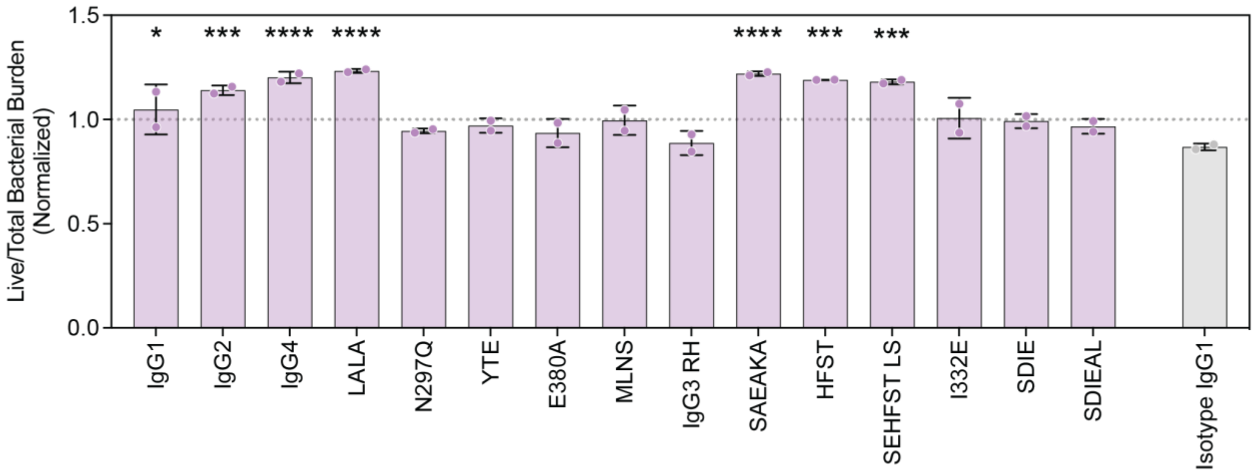


**Figure S2: Macrophage *Mtb* restriction assay of Fc-engineered α-glucan antibodies**. Y-axis shows live (GFP) / total (mCherry) *Mtb* burden in human monocyte-derived macrophages normalized by the no antibody condition for the respective donor. Each point is the triplicate average from 1 human macrophage donor. One-way ANOVA with Dunnett's correction comparing each antibody with the isotype IgG1 control antibody Adjusted p-value: < 0.05 (*), < 0.01 (**), < 0.001 (***), < 0.0001 (****). Error bars indicate mean with standard deviation.

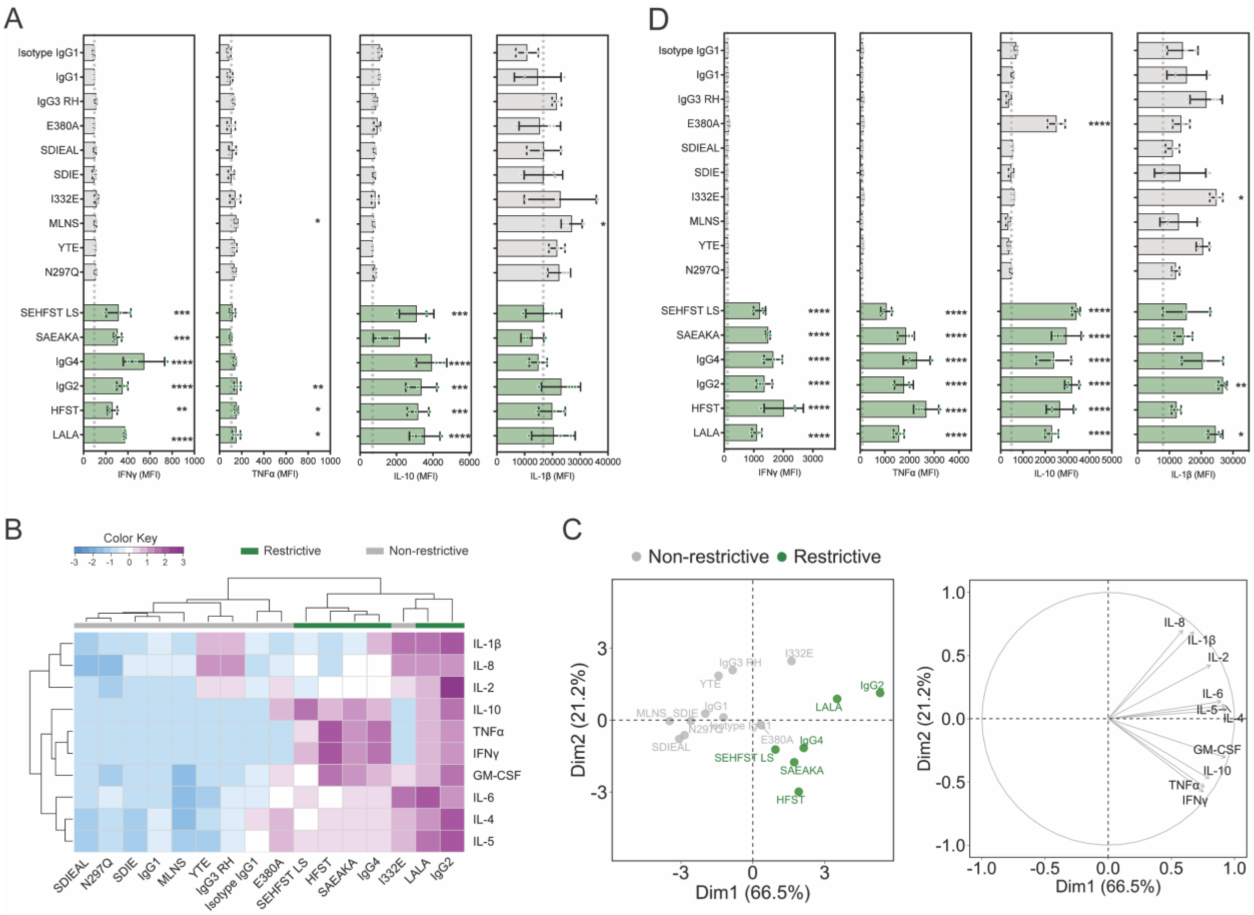

**Figure S3: Cytokine Luminex of whole-blood restriction assay.** Cytokine Luminex using the whole-blood assay supernatant collected at 120 hours. (**A** and **D**) Luminex MFI across selected cytokines in **(A)** Donor A, and **(D)** Donor B. One-way ANOVA with Dunnett's correction comparing each antibody with the isotype IgG1 control antibody. Green (Restrictive); grey (Non-restrictive). Adjusted p-value: < 0.05 (*), < 0.01 (**), < 0.001 (***), < 0.0001 (****). Error bars indicate mean with standard deviation. Run in technical triplicate. **(B)** Clustered heatmap indicating the cytokine profile elicited by each α-glucan Fc-variant in Donor B. Data were z-scored prior to heatmap visualization. (**C**) Principal component analysis of cytokine Luminex data from Donor B. Left, score plot of the first two principal components. Right, loading plot of the first two principal components.

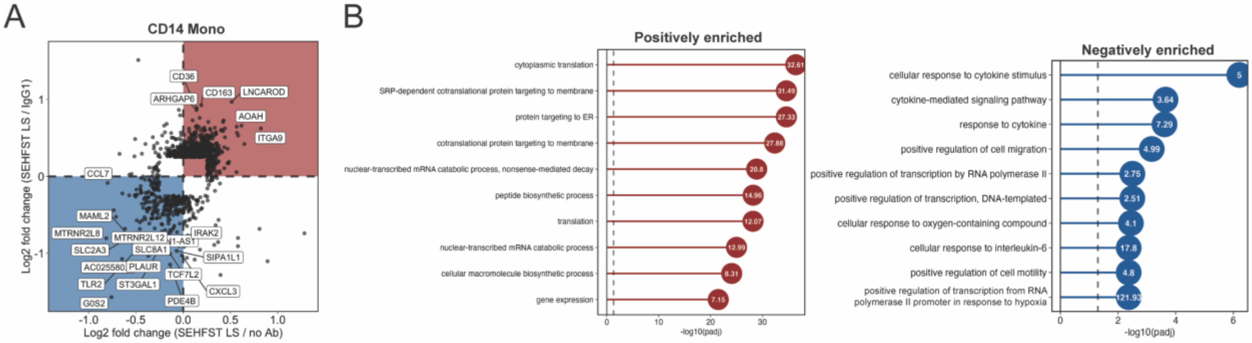


**Figure S4: CD14 monocyte differential expression analysis.** scRNAseq analysis of CD14 monocytes. **(A)** Genes consistently increased in the 24c5 SEHFST LS condition (red quadrant): (i) Mann-Whitney p-value < 0.1 and a log2 fold change > 0.25 compared to either the 24c5 IgG1 or no Ab condition, (ii) detected in a minimum fraction of 0.1 cells in either of the two conditions, and (iii) a log2 fold change > 0 compared to both the 24c5 IgG1 and no Ab conditions. Genes consistently decreased in the 24c5 SEHFST LS condition (blue quadrant): (i) Mann-Whitney p-value < 0.1 and a log2 fold change < -0.25 compared to either the 24c5 IgG1 or no Ab conditions, (ii) detected in a minimum fraction of 0.1 cells in either of the two conditions, and (iii) a log2 fold change < 0 compared to both the 24c5 IgG1 and no Ab conditions. **(B)** Gene list enrichment analysis using GO Biological Process gene sets. Left, GO terms enriched in red quadrant genes from panel C. Right, GO terms enriched in blue quadrant genes from panel C. Vertical dashed line indicates adjusted p-value of 0.05. Numbers on each circle show the odds ratio. Top ten GO terms by adjusted p-value shown.
